# NGAL deficiency elicits Hemophilia-like bleeding and clotting disorder

**DOI:** 10.1101/2023.09.15.557008

**Authors:** Min Xue, Shaoying Wang, Changjiang Li, Yuewei Wang, Ming Liu, Dandan Xiao, Qikai Yin, Liyuan Niu, Chuanbin Shen, Jianxun Wang, Xiaopeng Tang

## Abstract

Coagulation is related to inflammation, but the key pathways, especially innate immunity inflammatory response-coagulation, hemostasis, and thrombosis regulation is poorly understood and need to be further explored. In the current study, we showed that innate immunity inflammatory mediator neutrophil gelatinase-associated apolipoprotein (NGAL) which was upregulated in plasma of deep vein thrombosis patients interacted with and potentiated thrombin, kallikrein, FXIa, and FVIIa and suppressed antithrombin to induce coagulation, hemostasis, and thrombosis. Furthermore, NGAL can augment thrombin-induced platelet aggregation. In multiple mice hemostasis and thrombosis models, NGAL overexpression or intravenous administration promoted coagulation and hemostasis and aggravated thrombus, whereas NGAL knockout or treatment with anti-NGAL monoclonal antibody significantly prolonged bleeding time and alleviated thrombus formation. Notably, NGAL knockout prolonged both mice tail bleeding time and artery occlusion time to over 40 min, resembling uncontrollable bleeding and clotting disorder seen in Hemophilia mice. Furthermore, anti-NGAL monoclonal antibody treatment markedly reduced the formation of blood clots in a mouse-tail thrombosis model induced by carrageenan, which is linked to inflammation. Collectively, these findings suggest NGAL is a crucial coagulation regulator and mediates the crosstalk between innate immunity inflammation and coagulation, hemostasis, and thrombus, and provide new target and strategy for the development of innovative antithrombotic drugs.

## Introduction

Inflammation and coagulation regulation are closely related^1-5^. Studies have shown that tissue factor activates clotting factors, triggering pro-inflammatory responses, and initiating downstream cell signaling pathways^1,6-8^. Tissue factor is located at the intersection of coagulation and inflammation, serving as a trigger for host responses to injury or invasion by pathogens. Additionally, white blood cells, platelets, and endothelial cells release inflammatory cytokines and procoagulant microparticles (MPs), which sustain the exposure of tissue factor in the blood^1,3^. Thrombin plays a central role in controlling bleeding and thrombus formation, and exerts hemostatic and procoagulant effects through the coagulation-inflammation axis^1,7,9,10^. Thrombin can activate various types of cells by hydrolyzing protease-activated receptors (PARs)^11-13^, which are expressed in multiple cells, including platelets, immune cells, endothelial cells, and neurons^14^. After thrombin-mediated PAR activation, platelets degranulate, change shape, and aggregate. In terms of inflammation, thrombin activates PAR-1 in multiple cells, triggering the production of chemotactic protein-1, TNFα, and IL-6. PAR-1-dependent signaling also leads to endothelial cell activation, resulting in increased exposure of P-selectin and expression of monocyte chemotactic protein-1, plasminogen activator inhibitor-1, and IκB-β^1^. These actions of endothelial cells facilitate the adherence and recruitment of platelets and leukocytes, and play an important role in the early stages of venous thrombus formation^1^. Overall, these processes form a strong positive feedback loop, amplifying inflammation, and coagulation processes^1^. Furthermore, activated protein C (APC) inhibits the coagulation reaction by cleaving thrombin factors Va and VIIIa^3,6,7,10^. The APC-thrombomodulin-protein C receptor axis has significant vascular protective properties. APC induces PAR-mediated anti-inflammatory/cellular protective signals through endothelial protein C receptor, upregulates apoptosis inhibitors (IAP-1, A20) and sphingosine-1 phosphate (S1P) signaling pathways, and inhibits endothelial cell cytokine release. In addition, the contact activation of the coagulation system releases bradykinin (BK) ^15,16^. BK binds to B2R receptors on neutrophils and endothelial cells triggers a proinflammatory response.

Innate immune system regulates coagulation, hemostasis, and thrombosis^1,17-21^. Toll-like receptors and complement system of innate immunity related to inflammation play important roles in the pathogenesis of thrombosis^1-3,22^. Toll-like receptors are activated by lipopolysaccharide, DNA, heat shock proteins, etc., and produce inflammatory cytokines such as interferon, TNF-α, and IL-1β, thereby promoting tissue factor expression and coagulation. The complement system is an important inducer of inflammation and thrombosis^23,24^, and the binding of complement factors C3a and C5a to receptors C3aR and C5aR can mediate the early stages of inflammation and thrombosis^1,25^. C3a and C5a can also induce endothelial cells to express IL-8. The exposure of P-selectin on endothelial cells can be triggered by C5a, which plays a crucial role in facilitating the adhesion of neutrophils to endothelial cells^1,7,25^. Furthermore, C5a mediates tissue factor and hemophilia factor expression in neutrophils and endothelial cells. Monocyte tissue factor expression induced by C5b-7 and platelet activation mediated by C5b-8/C5b-9 can directly induce the formation of procoagulant phenotypes^1,7,25^. The fibrinolytic system is essential for the breakdown of blood clots, but interestingly, it can also contribute to inflammation processes associated with thrombosis^1,7^. For example, plasmin can directly activate the production of inflammatory C5a and C5b. Plasminogen can cleave iC3b, leading to the disruption of its ability to inhibit IL-12 expression in macrophages. Neutrophil extracellular traps (NETs) are formed by neutrophils and can capture circulating cells. The cellular components associated with NETs, such as histones, DNA, and proteases, have been implicated in inducing inflammation and potentially promoting the formation of blood clots^1,2,26-28^. Overall, our understanding of how the innate immune system regulates coagulation, hemostasis, and thrombosis is still limited, requiring further in-depth study^1^. Furthermore, an optimal therapy for the prevention or treatment of thromboembolism would aim to suppress excessive inflammation and coagulation without interfering with normal hemostasis or compromising the immune system^29^.

In the current study, we clarified that the innate immunity inflammatory mediator neutrophil gelatinase-associated apolipoprotein (NGAL) plays a significant role in the modulation of coagulation, hemostasis, and thrombosis and mediates the interaction between coagulation and inflammation. NGAL deficiency elicits uncontrollable bleeding and clotting disorder. It achieves this by interacting with and enhancing the activity of thrombin, kallikrein, FXIa, and FVIIa and inducing thrombin-induced platelet aggregation, while concurrently inhibiting antithrombin.

## Results

### NGAL is significantly upregulated in patients of deep vein thrombosis (DVT)

The hypercoagulable state, also known as the prothrombotic state, is characterized by an abnormality in the coagulation system that increases the risk of thrombosis^30^. Venous thromboembolism is the most common manifestation of this condition^30^. To investigate this further, we conducted an analysis of differential protein expression in the plasma of DVT patients and healthy individuals using data independent acquisition mass spectrometry (DIA-MS). As demonstrated in Figure 1A, we created a volcano plot based on the fold-change magnitude (log2 (DVT/Healthy)). Notably, NGAL (indicated by blue arrow) was found to be significantly increased in DVT patients. Additionally, we used an enzyme-linked immunosorbent assay (ELISA) to determine the average NGAL concentration in the plasma of DVT patients (n = 22), acute ST elevated myocardial infarction (STEMI) patients (n = 28), and healthy individuals (n = 36) (Table S1). Significantly, NGAL was markedly higher in DVT patients compared to STEMI patients, with levels measuring 322.2 ng/ml (SD 185.6) in DVT patients, 146.9 ng/ml (SD 52.1) in STEMI patients, and 66.2 ng/ml (SD 16.0) in healthy individuals (Figure 1B). These results suggest a potential link between NGAL and hypercoagulability.

**Figure 1.**
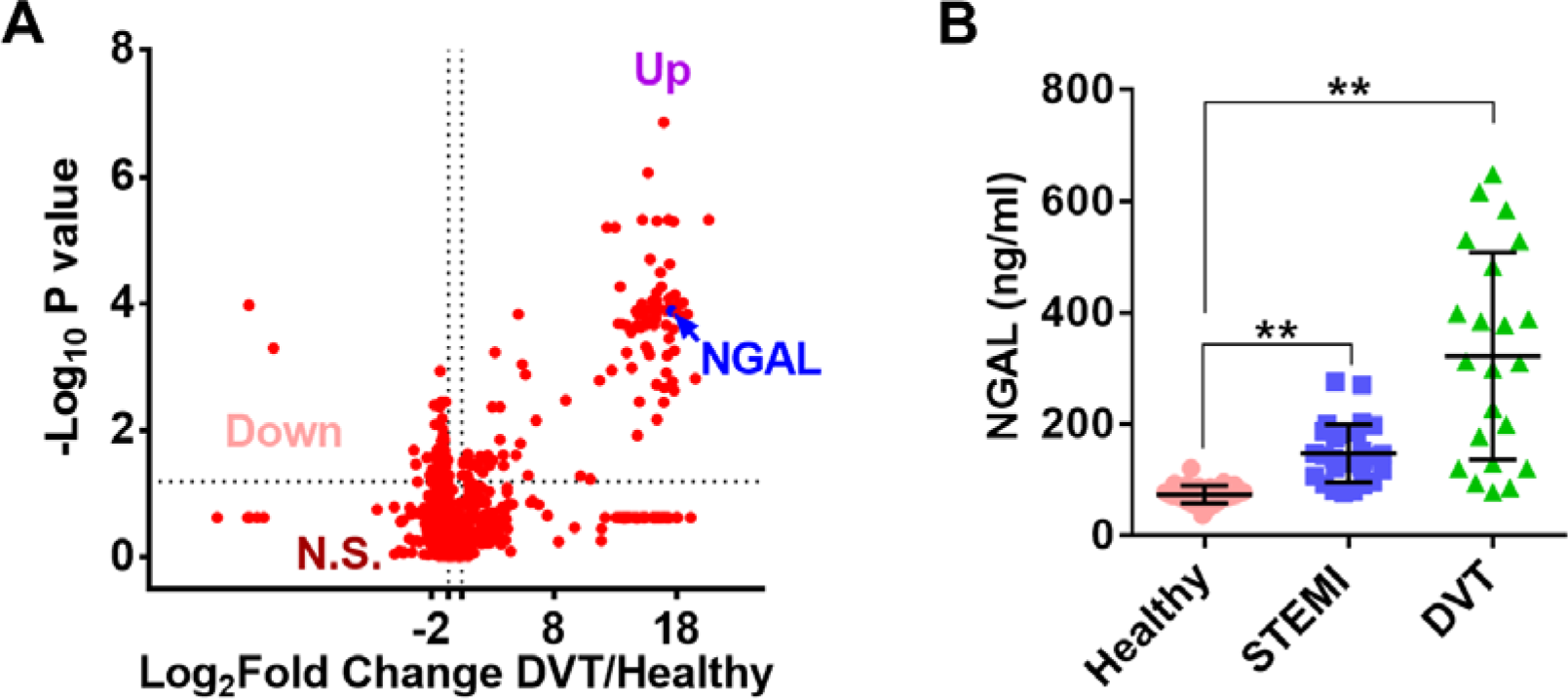
NGAL is significantly upregulated in plasma of deep vein thrombosis patients. **(A)** Volcano plot of all proteins quantified in plasma samples of deep vein thrombosis patients (DVT) patients (n = 3) and normal healthy volunteers (n = 3) from DIA-MS. Dots in top right corner represent up-regulated proteins in DVT with a significant fold-change (DVT/Normal); Dots in top left corner represent down-regulated proteins in DVT; Dots at below represent proteins with no obvious changes. NGAL which significantly upregulated in DVT was selected as a protein of interest for validation (indicated by blue arrow). **(B)** Amounts of NGAL in plasma from DVT patients, acute ST elevated myocardial infarction (STEMI) patients, and healthy volunteers were determined by ELISA. Data represent means ± SD (n=22-36), \*\**p* < 0.01 by Mann-Whitney U test. N.S.: no significance.

### NGAL potentiates multiple coagulation factors while inhibiting antithrombin

To investigate the potential mechanism underlying NGAL-induced hypercoagulability, various tests were conducted to assess the impact of NGAL on coagulation and platelet. As depicted in Figure 2A-C and Figure S1A and B, NGAL was found to promote coagulation by shortening the recalcification time without affecting platelet aggregation that was stimulated by ADP or collagen. The enzymatic activities of various coagulation factors and inhibitors were tested to assess the effects of NGAL on coagulation. Results showed that NGAL enhanced the activity of thrombin, kallikrein, FXIa, and FVIIa (Figure 1D and Figure S1C), while it had no impact on FXIIa, FIXa, FXa and plasmin (Figure 1E). Furthermore, NGAL also increased the enzymatic activity of thrombin, kallikrein, FXIa, and FVIIa towards their natural substrates: fibrinogen, FXII, and FIX (Figure 2F-I and Figure S1D-F), respectively. In Figure 2F, it can be observed that the levels of Fibrinopeptide A (FbpA) and FbpB, which are products of fibrinogen hydrolysis by thrombin, were enhanced approximately 1.2-, 2.3-, and 3.4-fold, and approximately 1.0-, 1.8-, and 3.0-fold, respectively, after treatment with NGAL at concentrations of 0.5, 1, and 2 μg/ml. At concentrations of 0.5, 1, and 2 μg/ml, NGAL enhanced kallikrein’s ability to release the hydrolytic product of FXII (FXIIa heavy chain (HC), ∼50 kDa) to approximately 1.2-, 1.4-, and 1.8-fold, respectively (Figure 2G and Figure S1D). As shown in Figure 1H and I, and Figure S1E and F, NGAL enhanced the capacity of both FXIa and FVIIa to hydrolyze FIX. This suggests that NGAL play a role in increasing the catalytic activity of FXIa and FVIIa towards FIX. In vitro studies showed that NGAL impeded the formation of thrombin-antithrombin (TAT) complexes and inhibits the activity of antithrombin against FIXa, as demonstrated in Figure 2J and K and Figure S1G. As illustrated in Figure 2L and Figure S1H, NGAL promoted thrombin-induced platelet aggregation. Thromboelastography analysis showed NGAL treatment strikingly decreased clot reaction time (R time, 2.3 vs 7.6 min) and clot kinetic time (K time, 1.5 vs 6.3 min) and increased thrombus generation angle (α angle, 63 vs 33 °) and thrombus maximum amplitude (MA value, 58.6 vs 52.3 mm), indicating its critical roles in inducing coagulation and platelet aggregation (Figure 2M). All these results suggest NGAL induces hypercoagulability and contributes to coagulation and hemostasis by enhancing multiple coagulation factors, neutralizing the inhibitory effects of antithrombin, and enhancing platelet aggregation.

**Figure 2.**
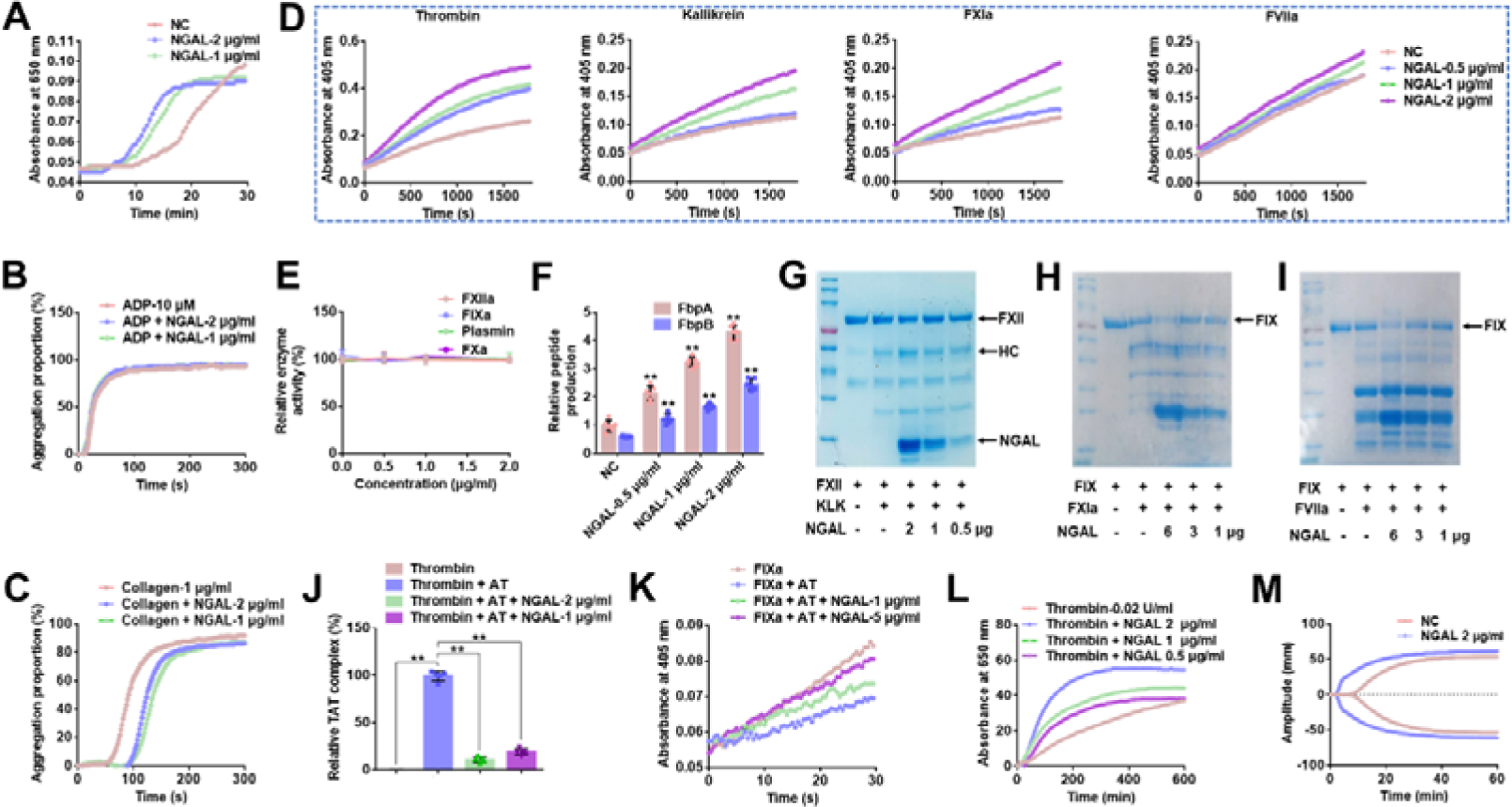
NGAL potentiates multiple clotting factors and inhibits antithrombin. **(A)** Plasma recalcification time was shortened by human NGAL. NGAL showed no effect on ADP (10 μM) **(B)** or collagen (1 μg/ml) **(C)** induced platelet aggregation. **(D)** Potentiating effects of NGAL on thrombin, kallikrein, FXIa, and FVIIa. **(E)** NGAL showed no influence on FXIIa, FIXa, FXa, and plasmin. **(F)** Representative ELISA quantification of fibrinopeptide A (FbpA) and FbpB released from fibrinogen hydrolyzed by thrombin mixed with 0.5, 1, or 2 μg/ml NGAL, respectively. **(G)** Representative SDS-PAGE analysis of FXII hydrolysis by kallikrein. Lane 1: 2 μg FXII; lane 2: 2 μg FXII + kallikrein; lane 3, 4, 5: 2 μg FXII + kallikrein + 2, 1, 0.5 μg/ml NGAL, respectively. Representative SDS-PAGE analysis of FIX hydrolysis by FXIa **(H)** or FVIIa **(I)**. **(J)** NGAL inhibits thrombin–antithrombin complex (TAT) formation by ELISA analysis. **(K)** NGAL blocked antithrombin (AT)’s inactivation effect on FIXa. **(L)** NGAL augmented thrombin-induced platelet aggregation. Effect of NGAL on Thrombelastography was shown. Data represent mean ± SD of 3-6 independent experiments, **p < 0.01 by one-way ANOVA with Dunnett’s post hoc test (F and J).

### NGAL directly interacts with thrombin, kallikrein, FXIa, FVIIa and antithrombin

As depicted in Figure 3A-E, results obtained from surface plasmon resonance (SPR) analysis indicated that NGAL exhibited direct interaction with thrombin, kallikrein, FXIa, FVIIa, and antithrombin. Our study found that NGAL and thrombin, kallikrein, FXIa, FVIIa or antithrombin showed interaction *in vitro*. Thus, we investigated the physiological occurrence of the NGAL-thrombin/kallikrein/FXIa/FVIIa/antithrombin interaction. Co-immunoprecipitation analysis revealed the physiological interaction between NGAL and prothrombin/prekallikrein/FXI/FVII/antithrombin in plasma (Figure 3F). As demonstrated in Figure 3G, confocal analysis also revealed the physiological interaction between NGAL and thrombin/kallikrein/FXIa/FVIIa/antithrombin in DVT. The presence of NGAL-thrombin-/kallikrein-/FXIa-/FVIIa-/antithrombin-positive deposits indicated the formation of NGAL-thrombin-/kallikrein-/FXIa-/FVIIa-/antithrombin complexes in thrombus (Figure 3G).

**Figure 3.**
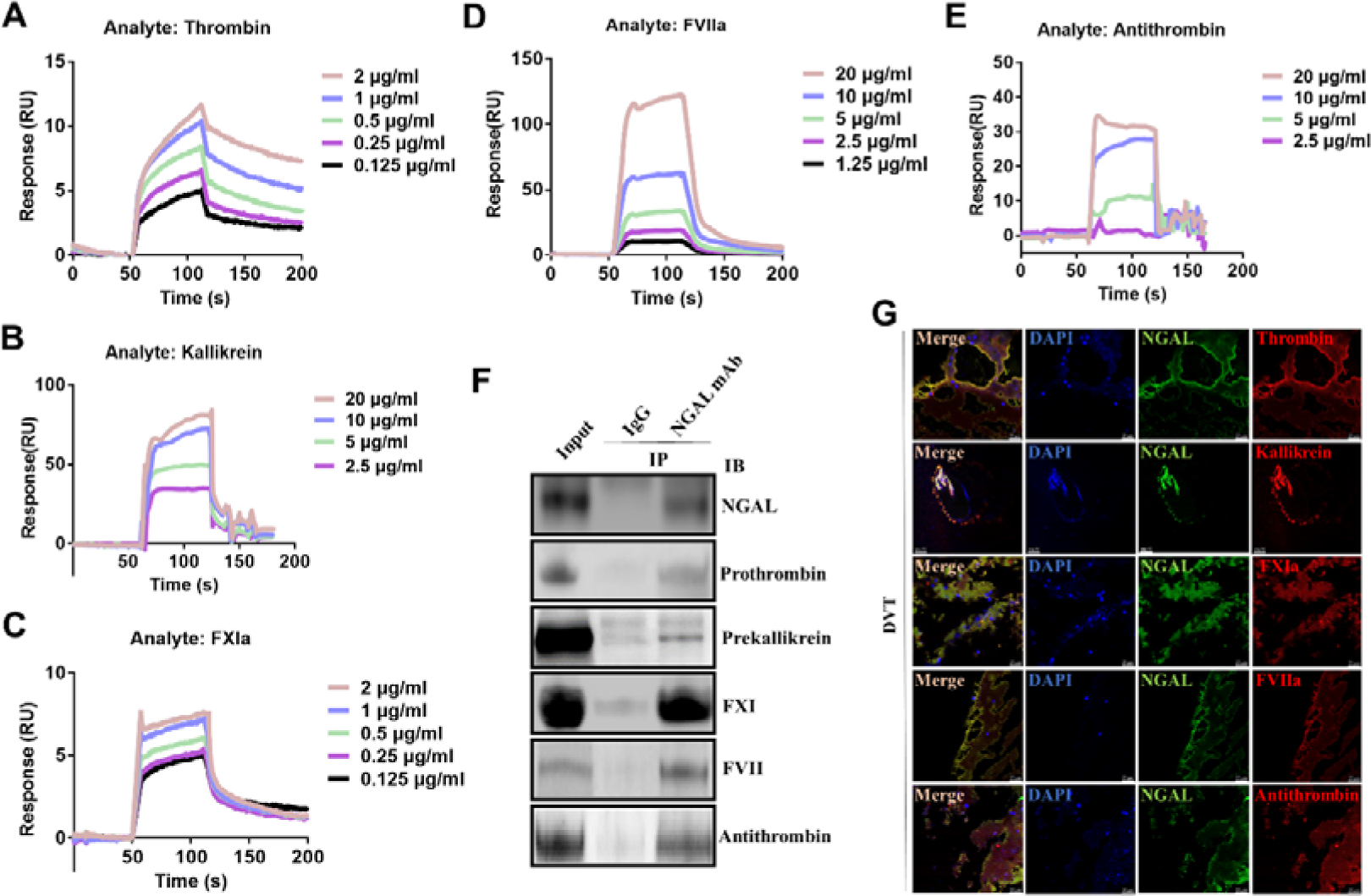
Direct interactions between NGAL and clotting factors. SPR analysis of the interaction between NGAL and thrombin **(A)**, kallikrein **(B)**, FXIa **(C)**, FVIIa **(D)** or antithrombin **(E)**. **(F)** Co-immunoprecipitation of NGAL and prothrombin, prekallikrein, FXI, FVII, or antithrombin in normal plasma. **(G)** Confocal analysis of NGAL-thrombin/kallikrein/FXIa/FVIIa/antithrombin-positive structures. Clots from DVT patients were labeled with NGAL (green) and thrombin, kallikrein, FXIa, FVIIa, or antithrombin antibody (red) to detect the presence of NGAL-thrombin/kallikrein/FXIa/ FVIIa/antithrombin complex. Cell nuclei were labeled by DAPI. Scale bar represents 25 µm. Images were representative of at least 3 independent experiments.

### NGAL overexpression promotes coagulation and hemostasis that are inhibited due to NGAL deficiency

To further investigate the function of NGAL in coagulation, we conducted experiments to evaluate the effects of NGAL overexpression, intravenous administration, NGAL monoclonal antibody interference, and NGAL deficiency on various indexes such as plasma thrombin activity, activated partial thromboplastin time (APTT), prothrombin time (PT), and thromboelastography. NGAL expression levels were initially confirmed through multiple analysis, including ELISA, western blot, and PCR (Figure S2A-C). Following injection of 10^12^ vg adeno-associated virus, an elevation in NGAL plasma concentration was observed in mice overexpressing NGAL (AAV9-NGAL) compared to controls (NC) or mice injected with a blank virus (AVV9) without NGAL overexpression (Figure S2A). Conversely, a deletion of plasma NGAL was observed in NGAL knockout mice (KO^NGAL^) compared to wild type (WT) mice (Figure S2B and C). Notably, NGAL knock out showed no effect on plasma iron concentration (Figure S2D). Results showed that plasma thrombin activity was significantly increased in plasma from AAV9-NGAL mice and mice with intravenous NGAL administration, while APTT and PT was shortened (Figure 4A-C). Conversely, NGAL monoclonal antibody-treated mice (NGAL mAb) showed significantly reduced plasma thrombin activity, prolonged APTT, and PT (Figure 4A-C). As expected, the blank virus controls (with empty AAV9 vector) and control IgG showed no effects on plasma thrombin activity, APTT, and PT (Figure 4A-C). Compared with negative group, NGAL mAb treatment prolonged R time, K time and reduced clot angle, while NGAL overexpression had the opposite effects as shown in thromboelastography analysis (Figure 4D). The role of NGAL in hemostasis was further investigated using mice tail-bleeding and saphenous vein bleeding models. Significantly shorten tail-bleeding and saphenous vein bleeding time was found in AAV9-NGAL and intravenous NGAL administration mice, with AAV9 control mice showing no effects (Figure 4E and F). Conversely, NGAL monoclonal antibody-treated mice (Figure 4E and F) showed significantly prolonged tail-bleeding or saphenous vein bleeding time, whereas the control IgG showed no effect on the prolongation of bleeding time (Figure 4E and F). Notably, compared 4 min in control mice, the tail-bleeding time in mice treated with the anti-NGAL antibody exceeded 15 min, with some going beyond 40 min (Figure 4E and F).

**Figure 4.**
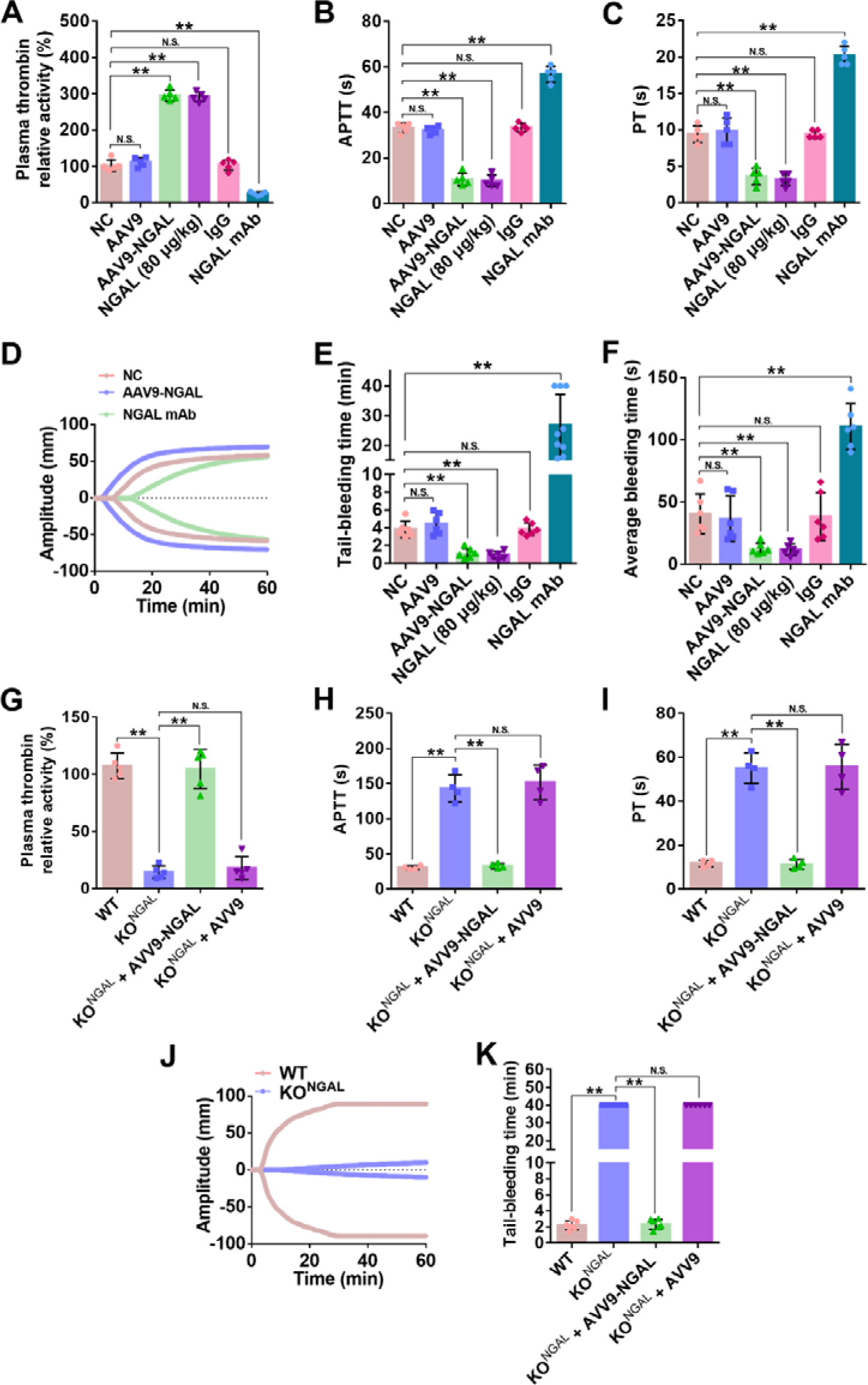
NGAL induces hypercoagulability and hemostasis which are reversed by NGAL deficiency. Effects of NGAL overexpression (AAV9-NGAL), blank virus (AAV9), intravenous injection of mice NGAL (80 μg/kg), anti-NGAL monoclonal antibody (NGAL mAB), or IgG control on plasma thrombin activity **(A)**, APTT **(B)**, PT **(C)**, thrombelastography **(D)**, mice tail-bleeding time **(E)**, and mice saphenous vein bleeding time **(F)**. **(G)** Effects of NGAL knock out (KO^NGAL^) on plasma thrombin activity **(G)**, APTT **(H)**, PT **(I)**, thrombelastography **(J)**, and mice tail-bleeding time **(K)**. Tail bleeding time was recorded for a maximum of 40 min to prevent mortality due to excessive bleeding and column 2 and 4 in panel “K” represent bleeding time that over 40 min. Data represent mean ± SD (n=4-9), **p < 0.01 by one-way ANOVA with Dunnett post hoc test or t-test. N.S.: no significance.

To further prove the critical role of NGAL on coagulation and hemostasis, NGAL knock out mice (Figure S2B and C) were performed to coagulation and hemostasis tests. As shown in Figure 4G-K, the knockout of NGAL significantly reduced plasma thrombin activity, MA value, and prolonged R time, K time, APTT, PT, and tail-bleeding time. The remarkably prolonged R time (13.4 vs 0.9 min), K time (83 vs 1.2 min), decreased clot angle (4.7 vs 68.7 °), and lower MA value (1.7 vs 82.2 mm) in NGAL knockout mice were consistent with reduced thrombin activity (14.4 vs 107.4 %), prolonged APTT (143.3 vs 30.5 s), and PT (55.0 vs 11.8 s) as depicted in Figure 4G-J. AAV9-NGAL overexpression which complement the NGAL deletion can reverse these changes in NGAL knock out mice, while AAV9 control has no effect (Figure 4G-K). Notably, tail-bleeding time in NGAL knock out mice were over 40 min when mice tail was cut 4.5 mm away with a bleeding phenotype like that of the hemophilia mice^31-35^. These findings collectively indicate that NGAL plays a crucial role in coagulation and hemostasis.

### Thrombus tendency is alleviated by NGAL deficiency, whereas NGAL overexpression has the opposite effect

The role of NGAL in hypercoagulability and thrombosis was further investigated using mice thrombosis model induced by FeCl_3_ and DVT model. Decreased artery blood flow (Figure 5A and B) and aggravated venous thrombus formation (Figure 5C and D) were observed in the carotid artery and deep vein of NGAL overexpression and intravenous NGAL administration mice, whereas NGAL monoclonal antibody-treated mice showed significant increased blood flow and alleviated venous thrombus formation (Figure 5A-D), with AAV9 control and IgG showing no effects (Figure 5A-D). To further investigate the role of NGAL in thrombosis, NGAL knock out mice was exerted in thrombus models described above. As illustrated in Figure 5E and F, at 10 % of FeCl_3_ induction, artery occlusion time in NGAL knock out mice was significantly prolonged and some over 60 min, whereas about 7.5 min in wild type mice. Furthermore, NGAL knock out can significantly inhibit venous thrombus formation (Figure 5G and H). NGAL complementation can reverse clotting disorder in NGAL knock out mice, while AAV9 control has no effect (Figure 5E-H). These results suggest that NGAL plays a key role in regulating thrombus pathogenesis.

**Figure 5.**
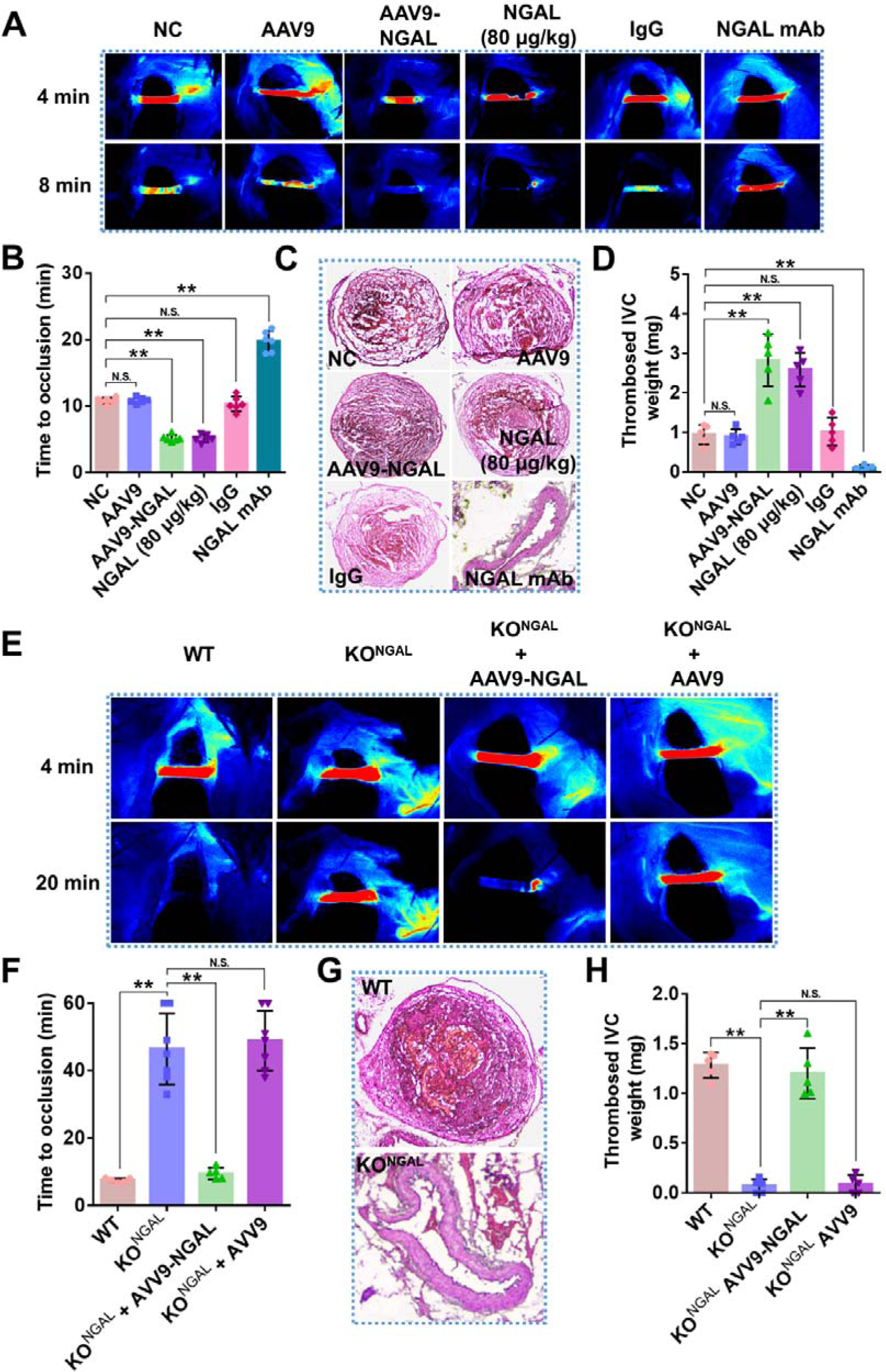
NGAL deficiency alleviates thrombus, which is aggravated by NGAL overexpression. Mice were received intravenous injection of blank virus (AAV9), NGAL overexpression virus (AAV9-NGAL), NGAL (80 μg/kg), anti-NGAL mAb (80 μg/kg), or IgG control (80 μg/kg). **(A)** Representative of carotid artery blood flow images in FeCl_3_-treated mice by laser speckle perfusion imaging, with region of interest placed in carotid artery to quantify blood flow change. **(B)** Time to occlusion was calculated in these mice group. Blood flow was recorded for a maximum of 60 min to prevent mortality. Mice were subject to inferior vena cava (IVC) stenosis for 24 hours to evaluate venous thrombogenesis. The pathological changes were observed through hematoxylin and eosin (HE) staining **(C)** and calculating thrombus weight **(D)**. Representative of carotid artery blood flow images of wild type (WT), NGAL knock out (KO^NGAL^), NGAL knock out with NGAL complementation by AAV9-NGAL (KO^NGAL^ + AAV9-NGAL) or blank AAV9 (KO^NGAL^ + AAV9) mice by laser speckle perfusion imaging were shown **(E)**, and time to occlusion was calculated **(F)**. HE staining **(G)** and thrombus weight **(H)** of IVC of mice described in “E” were shown. Data represent mean ± SD (n=5-8), **p < 0.01 by one-way ANOVA with Dunnett post hoc test or t-test. N.S.: no significance.

### NGAL deficiency alleviates carrageenan-induced mice tail thrombus

Thrombosis and inflammation are closely interconnected, and the administration of carrageenan in mice induced systemic inflammation, which subsequently led to the development of thrombosis in the tail vein^1,36-38^. Subsequently, we examined the impact of NGAL on carrageenan-induced tail thrombosis, which is associated with inflammation. After a 24 h induction, a substantial tail thrombus formation was observed in the carrageen-induced mice (Figure 6A-D). NGAL overexpression and intravenous administration increased tail vein thrombus, whereas NGAL monoclonal antibody-treatment can significantly reverse this tendency, with AAV9 control and IgG showing no effects (Figure 6A-D). Furthermore, NGAL knock out can significantly inhibit tail vein thrombus induced by carrageenan (Figure 6E and F). These results suggest that NGAL mediates the crosstalk between inflammation and coagulation and thrombus.

**Figure 6.**
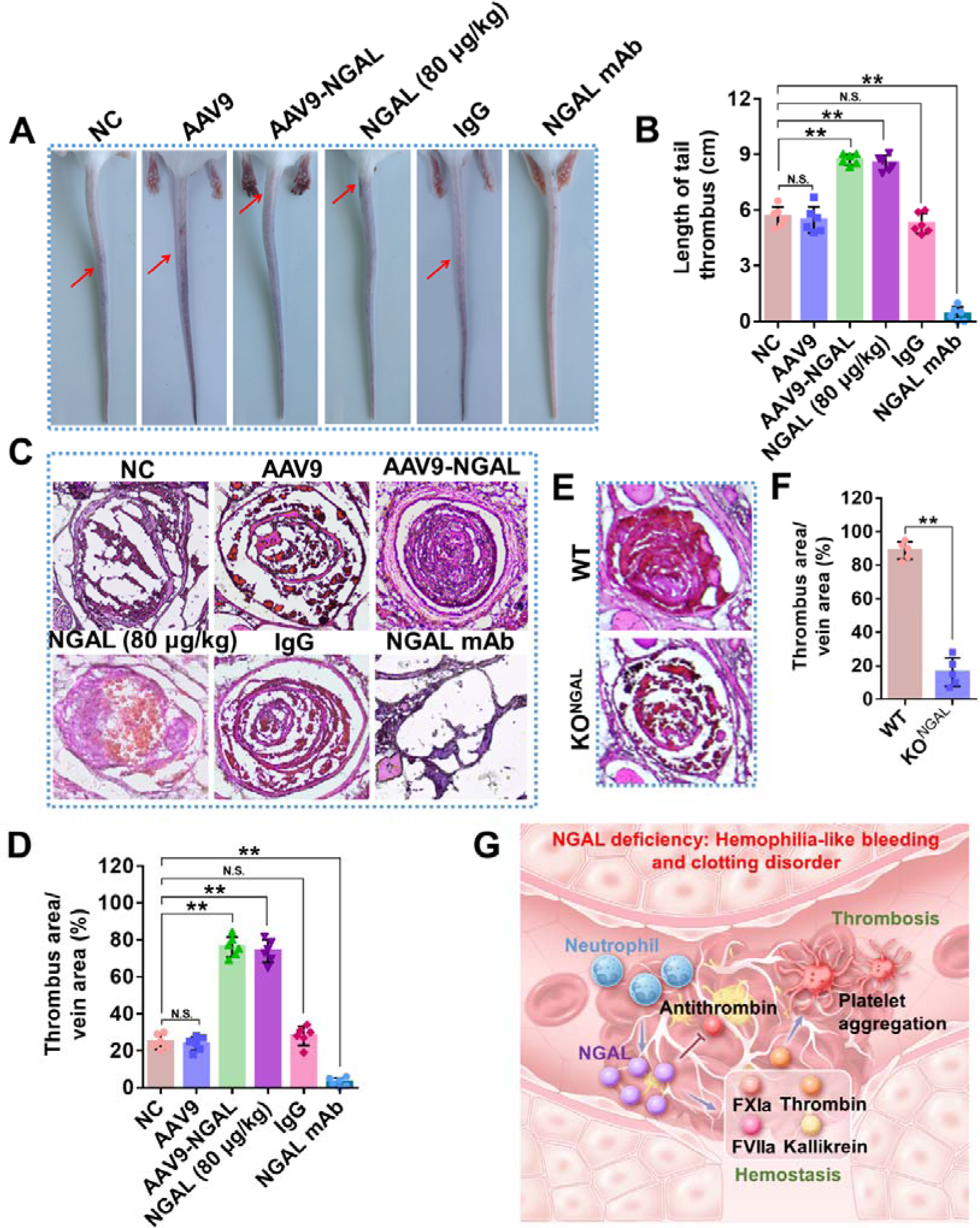
Carrageenan-induced mice tail thrombus is alleviated by NGAL deficiency, which is reversed by NGAL-supplementation. Mice were received intravenous injection of blank virus (AAV9), NGAL overexpression virus (AAV9-NGAL), NGAL (80 μg/kg), anti-NGAL mAb (80 μg/kg), and IgG control (80 μg/kg). **(A-B)** Thrombosis lengths in tails were measured and photographed at 24 h after carrageenan injection. **(C)** Hematoxylin and eosin (HE) staining of tail at 6 cm from the tail tip was shown. Quantification of “C” was shown on below **(D)**. HE staining of tail at 6 cm from the tail tip of wild type (WT) and NGAL knock out (KO^NGAL^) mice was shown **(E)**. Quantification of “E” was shown on right **(F). (G)** Graphic abstract of key role of NGAL in coagulation, hemostasis, and thrombosis and mediating crosstalk between coagulation and inflammation. NGAL knock out exerts Hemophilia-like bleeding and clotting disorder. Data represent mean ± SD (n=6), **p < .01 by one-way ANOVA with Dunnett post hoc test or t-test. N.S.: no significance.

## Discussion

Innate immunity is an important regulatory system for maintaining the internal environment stability of organisms, and excessive or insufficient strength can cause functional disorders in the body^39,40^. Neutrophils account for over 40% of all white blood cells and are the most abundant type of granulocyte, constituting the first line of defense in innate immunity and combating infections^41,42^. They are closely related to inflammation, infection, trauma, and cancer, and are natural candidates for performing important medical tasks in the human body^41^. Blood clots are rich in white blood cells, red blood cells, platelets, and fibrin. During the process of thrombosis and hemostasis, neutrophils are recruited to the site of clot formation and vascular damage by locally exposed and released von Willebrand factor, P-selectin, chemokines, and other factors, and may be involved in the pathological process of thrombosis and vascular injury^1,42,43^. Inflammatory reactions in neutrophils exhibit significant polarization, and platelets not only trigger the generation of NETs by physical contact but also affect the migration characteristics of neutrophils. In the pathological processes of sepsis and malignant tumors, NETs may be a major inducing factor for immune thrombosis^1,2,44^. It should be noted that the inflammatory response of neutrophils in the process of clot formation and vascular damage is a progressive process, including the release of various enzymes and active substances such as neutrophil elastase, suggesting the existence of important factors related to coagulation, hemostasis, and thrombus regulation that have not been identified^1,2^. The current study clarifies innate immunity inflammatory factor-NGAL plays critical roles in coagulation, hemostasis, and thrombus by potentiating multiple coagulation factors, augmenting thrombin-induced platelet aggregation, and inhibiting antithrombin and mediates the crosstalk between innate immunity and coagulation.

Previous reports have shown that NGAL is diagnosed urgently one of the most effective biological markers of sexual kidney injury and early diabetic nephropathy ^45,46^. NGAL is found from activated neutrophils in response to inflammation related to a relatively small molecular weight secretory protein^46^. As the important member of apolipoprotein family, NGAL is associated with inflammation, embryonic development, angiogenesis, diabetes, tumorigenesis, and atherosclerotic plaque instability^46-48^. NGAL has been reported to be associated with cardiovascular disease, with studies focusing on the correlation between NGAL and atherosclerotic plaque instability, myocardial damage, and myocardial fibrosis^46,49^. Research indicates that NGAL plays a role in the pathological mechanisms of stroke, such as glial activation, neutrophil infiltration, disruption of the blood-brain barrier, white matter injury, and neuronal cell death^49^. The current study showed NGAL deficiency in mice exerts Hemophilia-like bleeding and clotting disorder and unveiled a previously undisclosed role of NGAL in the processes of coagulation, hemostasis, and thrombosis.

It has been demonstrated NGAL can bind to proinflammatory ligands, including bacterial formyl peptides, leukotriene B4, and platelet-activating factor*^50,51^*. The current study clarified NGAL is a potentiator of thrombin and other multiple coagulation factors and augment thrombin-induced platelet aggregation without affecting ADP- or collagen-induced platelet aggregation. Collectively, NGAL plays a pivotal role in regulating hemostasis and thrombosis by exerting influence on both the coagulation cascade and platelet function, as also demonstrated in thrombelastography analysis. In multiple mice hemostasis and thrombus models, NGAL showed strong effect in influencing hemostasis and thrombus as a result. Our study found that NGAL knockout or treatment with monoclonal antibodies significantly extended the tail bleeding time in mice to over 40 min, resembling bleeding disorder observed in hemophilia mice in other documented reports^33,52,53^. Additionally, the knockout of NGAL and treatment with monoclonal antibodies notably extended the time of artery occlusion to over 60 min and 18 min, respectively, when 10 % FeCl_3_ was used as the inducer. The bacteriostatic properties of NGAL play a vital role in the antibacterial innate immune response by sequestering bacterial ferric siderophores, which results in iron depletion^54,55^. Our result indicates that the knockout of NGAL has no impact on plasma iron levels, suggesting that NGAL’s coagulation phenotypes are not dependent on iron. In addition, novel inhibitory peptides that interfere with the NGAL– coagulation factors interaction will be designed and studied using a docking model and analysis of their structural characteristics in future work. This research aims to develop a new strategy for the development of clinical anti-thrombus treatment, building on the approaches outlined in our previous studies^56-58^.

This discovery collectively elucidates the significance of NGAL as a crucial regulator of coagulation and highlights its novel role in facilitating communication between innate immunity, inflammation, hemostasis, and thrombosis (Figure 6G). NGAL deficiency exerts Hemophilia-like bleeding and clotting disorder, and offer a novel target and approach for effective antithrombotic medications (Figure 6G).

## Materials and Method

### Specimens of human plasma and thrombus sample of DVT

Samples of human plasma and thrombus were gathered from patients with DVT and STEMI with their informed consent prior to the study. The Institutional Review Board of the Qingdao University, the Affiliated Hospital of Qingdao University, and Qingdao Central Hospital of University of Health and Rehabilitation Sciences (QYFY WZLL 28251 and KY202319301) approved the study. Plasma specimens were collected from both DVT patients (n = 22, diagnosed through painful swollen lower limb and color duplex ultrasound combined with CT venography), STEMI patients (n = 28, diagnosed through chest pain accompanied by elevation of the ST segment in electrocardiogram), and healthy controls (n = 36) from Hospital (Table S1). Anticoagulant agent 1.5% EDTA-Na_2_ was used during the blood drawing process, and plasma was obtained via centrifugation at 3000 rpm for 20 min at 4 °C. The samples were sub-packed and then stored at -80 °C.

Thrombus specimens from 5 individuals of DVT were acquired through mechanical thrombectomy (Table S1) from Hospital. Immediately after surgical removal, some specimens were minced and homogenized at 4 °C for protein extraction. These were then sub-packed and stored at -80 °C. Other specimens were fixed in 10% buffered formalin to prepare frozen sections, while some were placed in RNAlater (R0901-500ML, Sigma, USA) for RNA extraction.

### Animals and ethics statement

Animal experiments conducted in this study were approved by the Animal Care and Use Committee at Qingdao University (QDU-AEC-2023364) and were conducted in accordance with Guide for the Care and Use of Laboratory Animals. NGAL knock out mice (#024630, 8 weeks of age, C57BL/6J background) were obtained from Jackson Laboratory while BALB/c mice (8 weeks old) were from Vitalriver Experiment Animal Company (Beijing, China). NGAL knock out mice was verified by PCR and western blot (Figure S2B-C). Housing for both strains was maintained in pathogen-free environments.

### NGAL measurement in plasma of DVT and STEMI

A human NGAL ELISA kit (SEB388Hu, Cloud-Clone Corp, China) was utilized to determine the concentration of NGAL in the plasma of DVT patients, STEMI patients, and healthy controls, following the manufacturer’s instructions.

### Effects of NGAL on coagulation and platelet aggregation

Plasma from healthy human donors was collected and prepared by mixing trisodium citrate (0.13 M) and blood in a 1:9 ratio. The mixture was then centrifuged at 3000 rpm/min for 30 min at 4 °C to obtain plasma. To study the effects of NGAL on plasma recalcification time, 20 μl of plasma was incubated with human NGAL (1 or 2 μg/ml, HY-P71156, MCE, USA) in 60 μl of HEPES buffer (20 mM HEPES, 150 mM NaCl, pH 7.4) for 10 min at 37 °C. This was followed by the addition of 60 μl of 25 mM CaCl_2_ preheated at 37 °C, and clotting was monitored at 650 nm using an enzyme-labeled instrument (Readmax 1500, Shanpu, Beijing, China). The time to half maximal increase in absorbance was measured to calculate clotting time.

Sodium citrate (1:9) was used as an anticoagulant agent during blood collection. Immediately after the blood draw, plasma of healthy was obtained by centrifugation at 150 g at room temperature for 10 min to separate platelet-rich plasma. NGAL (1 or 2 μg/ml) was then added to the platelets, which were incubated at 37 °C for 5 min. Collagen (1 μg/ml) or ADP (10 μM) was used to elicit platelet aggregations, which were detected by a platelet aggregator (AG400, Techlink biomedical, China). The maximum platelet aggregation rate was analyzed. Furthermore, platelet-rich plasma was centrifuged at 400 g for 10 min at room temperature to obtain platelet pellet. Platelets were washed in Tyrode’s buffer A (137 mM NaCl, 2 mM KCl, 0.3 mM NaH_2_PO_4_, 12 mM NaHCO_3_, 5.5 mM glucose, 1 mM MgCl_2_, 0.2 mM EDTA, and 0.35% BSA; pH 6.5) and then resuspended and in Tyrode’s buffer B (137 mM NaCl, 2 mM KCl, 0.3 mM NaH_2_PO_4_, 12 mM NaHCO_3_, 5.5 mM glucose, and 0.35% BSA; pH 7.4). The resuspended platelets were diluted to 2×10^8^/ml and then incubated with NGAL at 37 °C for 5 min. Aggregations were elicited by thrombin (0.02 U/ml).

### Effects of NGAL on activity of coagulation factors or inhibitor

The effect of NGAL on coagulation-related enzymes and inhibitors such as kallikrein, FXIIa, FXIa, FXa, FIXa, FVIIa, thrombin, plasmin, or antithrombin was evaluated using chromogenic substrates correspondingly. Specifically, the enzyme to be tested was incubated with NGAL (0.5, 1, or 2 μg/ml) in a 60 μL Tris-HCl buffer (50 mM, pH 7.4) for 5 min. Then, a particular concentration of chromogenic substrate was added according to the descriptions provided below. The absorbance at 405 nm was immediately measured using an enzyme-labeled instrument (Readmax 1500, Shanpu, China) and a kinetic curve recorded. The relative enzyme activity was calculated by measuring the enzymatic hydrolysis velocity of its substrate. Human α-thrombin (20 nM, HT 1002a, Enzyme Research Laboratories, USA) and human plasmin (20 nM, HPlasmin, Enzyme Research Laboratories, USA) were incubated with 0.2 mM chromogenic substrate S-2238 (Chromogenix AB, Sweden). Human α-FXIIa (50 nM, HFXIIa 1212a, Enzyme Research Laboratories, USA) and kallikrein (20 nM, HPKa 1303, Enzyme Research Laboratory, USA) were incubated with 0.2 mM chromogenic substrates S-2302 (Chromogenix AB, Sweden). For FIXa (HFIXa 1080, Enzyme Research Laboratory, USA) and FVIIa (HFVIIa, Enzyme Research Laboratory, USA), a concentration of 20 nM and 100 nM, respectively, were used, with the corresponding chromogenic substrates being 0.2 mM S-2288 (Chromogenix AB, Sweden). For FXa (HFXa 1011, Enzyme Research Laboratory, USA), a concentration of 30 nM was used, with the corresponding chromogenic substrate being 0.2 mM S-2222 (Chromogenix AB, Sweden). Moreover, the concentration used for FXIa (HFVIIa, Enzyme Research Laboratory, USA) was 20 nM, with the chromogenic substrate being 0.2 mM S-2366 (Chromogenix AB, Sweden). For a brief overview of the experiment of NGAL on antithrombin activity, NGAL with a concentration range of 1, 2 or 5 μg/ml, along with antithrombin (2 μM, HAT, Enzyme Research Laboratory, USA) and thrombin (10 nM) or FIXa (20 nM), were incubated concurrently in 60 μl of Tris-HCl buffer (50 mM, pH 7.4) for 5 min at 37 °C. The thrombin or FIXa activity was then evaluated using corresponding chromogenic substrates, like the approach used for thrombin alone or incubated with antithrombin, also tested in the same buffer. The TAT complex was assessed using an ELISA kit (SEA831Hu 96T, Cloud-Clone Corp, China) per the manufacturer’s instructions.

Furthermore, the effect of NGAL on the ability of thrombin to hydrolyze its natural substrate, fibrinogen, was evaluated. α-thrombin (20 nM) was incubated with fibrinogen (10 mg/ml) in Tris-HCl buffer (25 mM, pH 7.4), containing 0.15 M NaCl, and NGAL (0.5, 1, or 2 μg/ml) for 30 min at 37 °C. The concentration of FbpA and FbpB in supernatant was analyzed by using FbpA and FbpB ELISA kits (CEB237Hu 96T and CEB307Hu 96T; Cloud-Clone Corp, China). The effect of NGAL on kallikrein or FXIa to hydrolyze its natural substrate FXII or FIX was assayed by SDS–PAGE. Briefly, FXII (2 μg, HFXII1212, Enzyme Research Laboratory, USA) or FIX (2 μg, HFXII1212, Enzyme Research Laboratory, USA), was incubated with human kallikrein or FXIa (80 or 100 nM) in 40 μL of Tris–HCl buffer (50 mM, pH 7.4) in the presence of NGAL. After 20 min of incubation at 37 °C, all reactions were applied to 12% SDS–PAGE. For effect of NGAL on FVIIa to hydrolyze its natural substrate FIX, FIX (2 μg) was incubated with human FVIIa (800 nM) in 40 μL of Tris–HCl buffer (50 mM, pH 7.4) with 25 mM CaCl_2_ in the presence of NGAL for 30 min of at 37 °C, and all reactions were applied to 12% SDS–PAGE. The hydrolyzation production was detected.

### SPR analysis

The SPR experimental method was previously described^56^. In summary, NGAL was initially diluted to a concentration of 20 μg/ml with 200 μl of 10 mM sodium acetate at pH 5, and then flowed across the activated surface of the CM5 sensor chip for coupling. The remaining activated sites on the CM5 sensor chip were blocked using 1 M ethanolamine at pH 8.5. Various concentrations of thrombin (0.125, 0.25, 0.5, 1, and 2 μg/ml), kallikrein (2.5, 5, 10, and 20 μg/ml), FXIa (0.125, 0.25, 0.5, 1, and 2 μg/ml), FVIIa (1.25, 2.5, 5, 10, and 20 μg/ml), or antithrombin (2.5, 5, 10, and 20 μg/ml) in 20 mM Tris-HCl buffer at pH 7.4 were applied to analyze interactions.

### Confocal microscopy

To perform immunofluorescence detection of NGAL-thrombin/kallikrein/FXIa/FVIIa/antithrombin complexes in the thrombus specimens of DVT, 8-μm sections were prepared by cutting the specimens using a freezing microtome. Next, the sections were incubated with anti-human NGAL (dilution 1:200, ab125075, abcam, USA), anti-thrombin (dilution 1:200, P00734, Abmart, China), anti-FXIa (dilution 1:200, MabHFXI, Enzyme Research Laboratory, USA), anti-FVIIa (dilution 1:200, MabHFVII, Enzyme Research Laboratory, USA), anti-human kallikrein (dilution 1:200, ab1006, Abcam, USA), or anti-antithrombin (dilution 1:200, MabHAT, Enzyme Research Laboratory, USA) antibody overnight at 4 °C. Thereafter, the excess primary antibody was removed by washing the sections three times with PBS, followed by incubation with a fluorescently labeled secondary antibody for 1 h at 37 °C. The cell nuclei were stained with 4, 6-diamidino-2-phenylindole (DAPI, P36941, Life technologies, USA). Finally, images of immunofluorescence were acquired using the Leica confocal microscope (STELLARIS 5), following the manufacturer’s instructions.

### Immunoprecipitation

Normal plasma (0.5 μl) was mixed with 2 μg of anti-NGAL antibody and incubated for 16 hours at 4 °C in 200 μl of PBS (0.01M, pH 7.4). Following this, 20 μl of Protein A agarose (P2006, Beyotime, China) was added and incubated for 3 hours at 4 °C. The mixture was then centrifuged at 2,500 rpm for 5 min at 4 °C and washed with PBST (PBS + 0.05% Tween 20) for 7 times. Subsequently, 10 μl of loading buffer (1 × 20315ES05, Yeasen Biotechnology, shanghai, China) was added, and the coupled proteins were boiled for 3 min. All proteins were separated using 12% SDS-PAGE and probed with antibodies against NGAL, thrombin, FXIa, FVIIa, kallikrein, or antithrombin described above for the identification of target proteins.

### Effects of NGAL deficiency and supplementation on coagulation, hemostasis, and thrombosis

Recombinant adeno-associated virus serotype 9 (AAV9) expressing NGAL and control virus were designed and manufactured by JiKai Gene Chemical Technology Co., Ltd (Shanghai, China), and 10^12^ vg virus was injected into the tail vein of mice to induce NGAL overexpression. Effects of NGAL overexpression, mice NGAL intravenous injection (80 μg/kg, HY-P70658A, MCE, USA), NGAL mAb (80 μg/kg, 50060-R211, Sino biological, China), IgG control (80 μg/kg, ab199376, Abcam, USA) or NGAL knockout on thrombin activity, APTT, PT, thromboelastography, bleeding, and thrombus were tested as methods described below.

### Enzymatic activity assays of thrombin in plasma

Thrombin fluorogenic substrate was used to test the enzyme activity, following the same method as before^56,58^. Basically, we added 20 μl of the substrate (0.8 mM, Z-Gly-Gly-Arg-AMC, Bachem, Switzerland) with 20 mM CaCl_2_ into a 96-well plate containing 80 μl of plasma mixed with 20 μl of HEPES buffer, 1 pM tissue factor and 4 μM phospholipids to start the reaction. We then measured the fluorescence every 30 seconds for 30 min using a microtiter plate fluorometer (SpectraMax, Molecular Devices, Austria). The excitation and emission wavelengths were 390 and 460 nm, respectively.

### APTT and PT assays

Following our previous studies^56-58^, to test the APTT, a 100 μl sample of plasma was incubated with 100 μl of APTT reagent (R41056-50T, Yuanye, China) for 5 min at 37 °C, followed by the addition of 100 μl of CaCl_2_ (25 mM) preheated at 37 °C to test clotting time. The PT assay kit (product number GMS10176 from Genmed Scientifics, USA) was used as the methods described in manufacturer’s instructions.

### Thromboelastography analysis

Whole process of coagulation from the aspects of coagulation, platelet aggregation, and fibrinolysis were dynamically detected using a thromboelastogram analyzer (CWPS-8800, PMDT, China). The formation of natural clots was evaluated by mixing 20 μL of 0.2 M CaCl_2_ and 340 μl of whole blood into a test cup and the following values, for example, R time, K time, α angle, and MA value were monitored.

### Bleeding time measurement

To measure the bleeding time, the tip of each mouse tail was precisely cut 4.5 mm away and submerged in 20 ml of 37 °C pre-warmed saline solution, following established protocols^56-58^, with slight modifications. The bleeding time within 40-min time was monitored to prevent mortality due to excessive bleeding or recorded until the flow of blood ceased.

### Saphenous vein bleeding model

The saphenous vein bleeding model was carried out in accordance with our previously published protocols^58^. In brief, the saphenous vein of the mice was exposed and partially transected, and the time for the initial arrest of bleeding was recorded. The thrombus was then removed to elicit rebleeding, and this process was repeated over a duration of 30 min. The average hemostatic time within this 30-min time frame was documented.

### FeCl_3_-induced carotid artery thrombosis

To induce carotid artery thrombosis, FeCl_3_ was utilized as previously outlined^56-58^, with slight modifications. The mice were put under anesthesia using isoflurane inhalation facilitated by an anesthesia respirator (R540IP, RWD Life Science, China. Thrombosis was induced in the carotid artery by applying a filter-paper disk soaked in 10 % FeCl_3_. Carotid blood flow was continuously monitored for 60 min or until complete occlusion occurred using laser speckle perfusion imaging (RFLSI III, RWD Life Science, China) post FeCl_3_ induction, with perfusion units utilized to quantify blood flow.

### Mice DVT model

The mice DVT model was established by inducing stasis blood flow via infrarenal inferior vena cava (IVC) ligation^58,59^. In brief, the mice were anesthetized and midline laparotomy was performed. The venous side and dorsal branches of the IVC were obstructed, and stasis thrombosis was induced through ligation of the infrarenal IVC using a 7-0 Prolene suture (Ethicon Inc, Somerville, NJ). The peritoneum was closed using 5-0 Vicryl suture (Ethicon Inc, Somerville, NJ). The pathological changes were examined through HE staining, and the weight of the thrombus was calculated 24 hours later.

### Mouse-tail thrombosis model induced by carrageen

This model was performed in accordance with our previously protocols^60^. The mice received an intraperitoneal injection of 1% Type I Carrageenan (60 mg/kg, 9062-07-1, Coolaber, China), and were then maintained at 16 °C. The thrombosis region lengths in the tails were measured and photographed at 24 h after the carrageenan injection. The pathological changes of mice tail were examined through HE staining.

### Statistical analysis

The results from independent experiments were presented as the mean ± SD. All statistical analyses were two-tailed and with 95% confidence intervals (CI). Normal distribution was assessed using the Kolmogorov-Smirnov test (K-S test), followed by one-way ANOVA with post hoc Dunnett for p values. Unpaired t-test was utilized for comparisons between two groups. Nonparametric data were analyzed using the Mann-Whitney U test. Prism 6 (Graphpad Software) and SPSS (SPSS Inc, USA) were employed for data analysis. Statistical significance was considered at p < 0.05.

## Acknowledgments

This work was supported by the National Science Foundation of China (32371162), Excellent Young Scientists Foundation of Shandong Province (ZR2023YQ025), Taishan Scholars Program for Young Experts of Shandong Province, Start-up funding from Qingdao University (DC2300000302), and Natural Science Foundation of Shandong Province (ZR2019ZD28 and JQ20181).

## Author contributions

M.X., S.W., C.L., Y.W., M.L., D.X., and Q.Y. performed the experiments and data analyses; C.L., Y.W., and L.N. collected plasma and thrombus sample of DVT; X.T., J.W., and C.S. conceived and supervised the project; X.T. prepared the manuscript. All authors contributed to the discussions.

