## Supplemental methods, Figure 1-2, and Table 1 for "NGAL deficiency elicits Hemophilia-like bleeding and clotting disorder"

**Supplementary information**

**Supplementary methods**

**Data Independent Acquisition Mass Spectrometry (DIA-MS)**

Plasma samples were obtained from individuals diagnosed with deep vein thrombosis (DVT, n= 3) and healthy individuals (n= 3). DIA-MS was performed at Genefund Biotech Co., Ltd. (Shanghai, China) to identity the differentially expressed proteins in DVT.

**PCR analysis for NGAL deficient mice**

DNA was isolated from the toes of mice by using Quick Genotyping Assay Kit for Mouse Tail (D7283S, Beyotime, China) following manufacturer’s instruction. Primers 5’-AGCCAATGTAATCCAATCA and 5’-CCTAAGTCCCGTTCAATC amplify a region in exon 2 to exon 5 of NGAL gene (Figure S2B-a). Primers 5’-TAGGGGATGCCACATCTCA and 5’-TGGAGGTGACATTGTAGCTATTG were used to identify the NGAL gene (Figure S2B-b) while primers 5’-TAGGGGATGCCACATCTCA and 5’-CCTTCTATCGCCTTCTTGACG amplify a region within a neo cassette in NGAL deficient mice (Figure S2B-c) according to the suggestions from the Jackson Laboratory.

**Supplementary Table**

**Table S1. Clinical features and plasma NGAL concentrations in patients with DVT, STEMI, and healthy controls.**

|  | DVT | STEMI | Control (normal physical examinees) |
| --- | --- | --- | --- |
| Gender (Male:Female) | 1.75:1 | 1.8:1 | 1.4:1 |
| Age (years) | 24–76; 57 (14.3) | 37–89; 65 (13.5) | 26–69; 50 (12.5) |
| Main clinical features | Painful swollen lower limb and Visible venous thromboembolism | Chest pain accompanied by elevation of the ST segment in electrocardiogram | NA |
| NGAL (ng/ml) | 322.2 (185.6) | 146.9 (52.1) | 66.2 (16.0) |

DVT: deep vein thrombosis; STEMI: acute ST elevated myocardial infarction; NA: not applicable. Data represent mean (SD).

**Supplementary Figures and Figure legends**

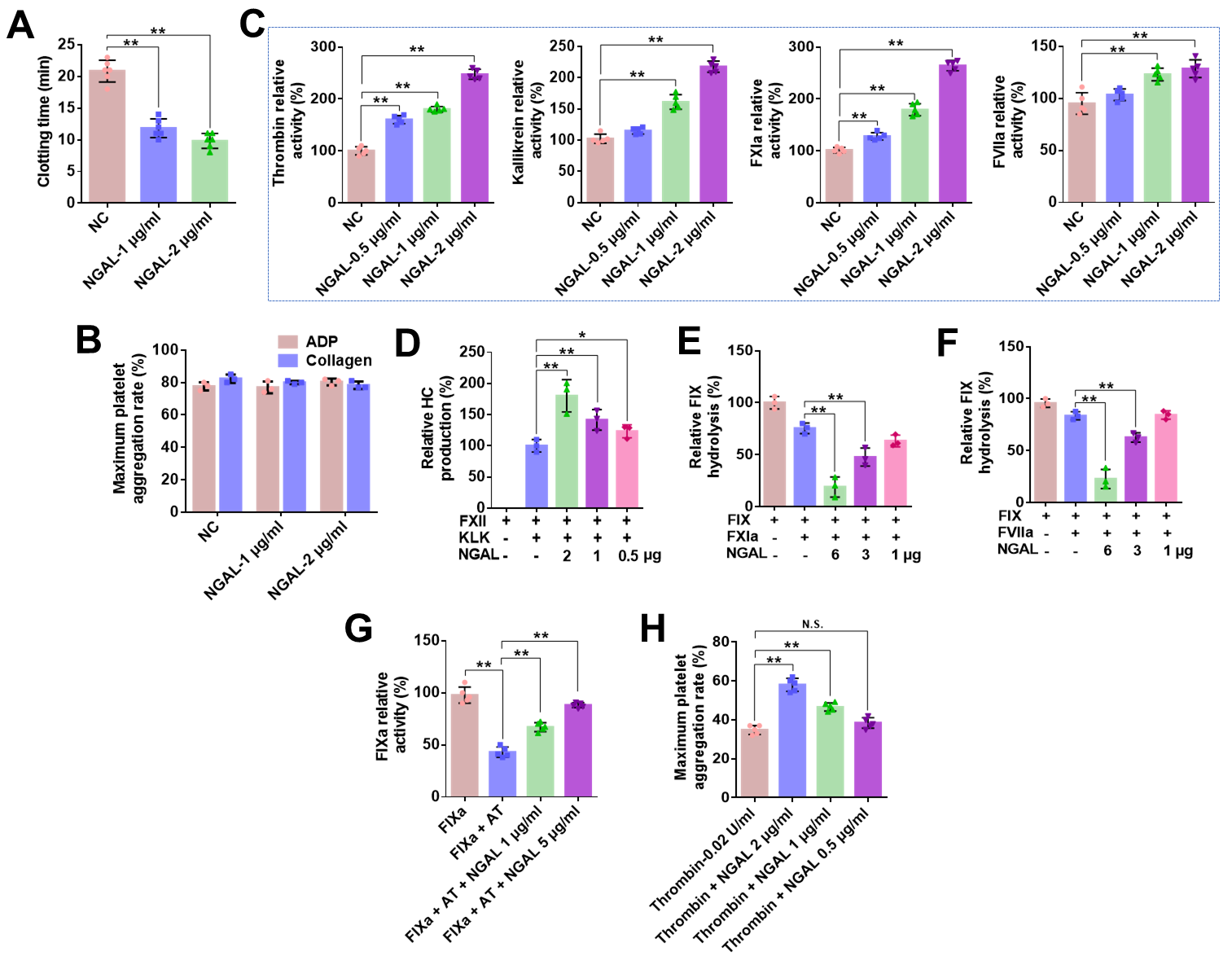

**Figure S1. Quantification of panels illustrated in Figure 2.** Data represent mean ± SD of 3-6 independent experiments, **p < 0.01 by one-way ANOVA with Dunnett’s post hoc test.

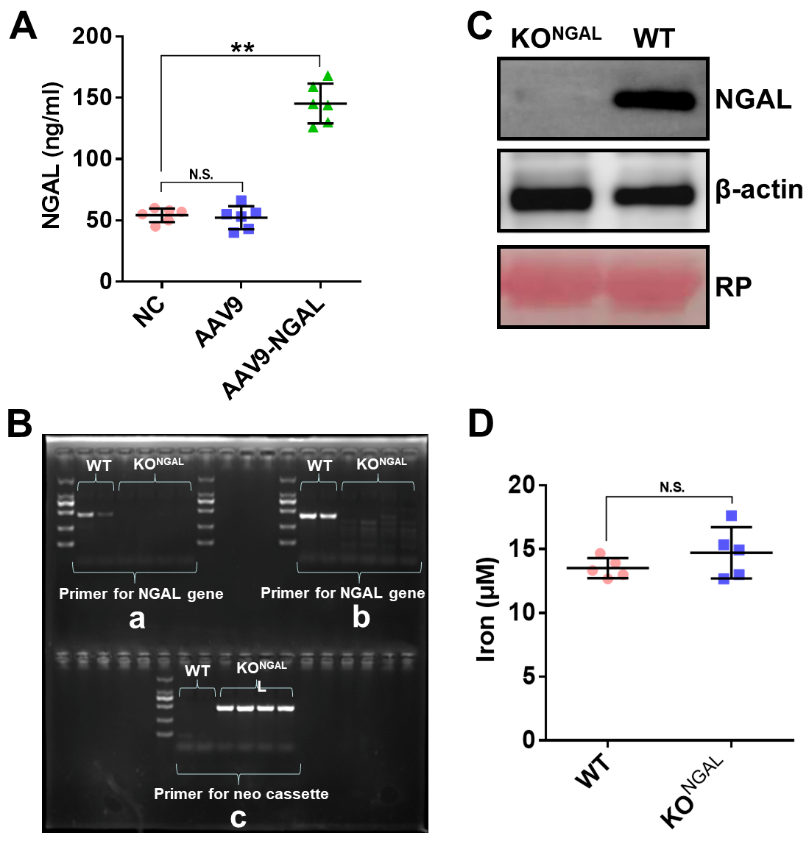

**Figure S2. Validation of the overexpression and knock out of NGAL.** Mice were received intravenous injection of blank virus (AAV9), NGAL overexpression virus (AAV9-NGAL). NGAL expression in plasma of these mice was determined by ELISA **(A)**. **(B)** PCR was performed to validate NGAL knock out mice. Primers 5’-AGCCAATGTAATCCAATCA and 5’-CCTAAGTCCCGTTCAATC amplify a region in exon 2 to exon 5 of NGAL gene (a); Primers 5’-TAGGGGATGCCACATCTCA and 5’-TGGAGGTGACATTGTAGCTATTG were used to identify the NGAL gene; Primers 5’-TAGGGGATGCCACATCTCA and 5’-CCTTCTATCGCCTTCTTGACG amplify a region within a neo cassette in NGAL deficient mice (c). **(C)** NGAL expression in plasma of wild-type (WT) and NGAL-knockout (KO^NGAL^) was determined by western blotting. **(D)** NGAL knockout had no effect on iron levels in plasma. Data are means ± SD (n = 8), ***p* < 0.01 by unpaired *t-*test. N.S.: no significant.
